# Experimental infection of Mexican free-tailed bats (*Tadarida brasiliensis)* with SARS-CoV-2

**DOI:** 10.1101/2022.07.18.500430

**Authors:** JS Hall, E Hofmeister, HS Ip, SW Nashold, AE Leon, CM Malavé, EA Falendysz, TE Rocke, M Carossino, U Balasuriya, S Knowles

## Abstract

The severe acute respiratory syndrome coronavirus-2 (SARS-CoV-2) virus originated in wild bats from Asia, and as the resulting pandemic continues into its third year, concerns have been raised that the virus will expand its host range and infect North American wildlife species, including bats. Mexican free-tailed bats (*Tadarida brasiliensis*: TABR) live in large colonies in the southern United States, often in urban areas, and as such, could be exposed to the virus from infected humans. We experimentally challenged wild TABR with SARS-CoV-2 to determine the susceptibility, reservoir potential, and population impacts of infection in this species. Of nine bats oronasally inoculated with SARS-CoV-2, five became infected and orally excreted moderate amounts of virus for up to 18 days post inoculation. These five subjects all seroconverted and cleared the virus before the end of the study with no obvious clinical signs of disease. We additionally found no evidence of viral transmission to uninoculated subjects. These results indicate that while TABR are susceptible to SARS-CoV-2 infection, infection of wild populations of TABR would not likely cause mortality. However, the transmission of SARS-CoV-2 from TABR to or from humans, or to other animal species, is a distinct possibility requiring further investigation to better define.

## Introduction

As we enter the third year of the severe acute respiratory syndrome coronavirus-2 (SARS-CoV-2) pandemic, many unanswered questions remain regarding the ecology of the virus. How the virus interacts with wild species is critical knowledge to obtain, including whether: (1) North American wildlife can act as reservoirs of the virus, (2) the virus can adapt genetically to new hosts and become more virulent, and (3) the virus can affect the health of wild populations, particularly threatened or endangered species.

SARS-CoV-2 naturally infects several wild species including captive wild animals. For example, American mink (*Neovison vison*) were infected with SARS-CoV-2 in the United States in proximity to domestic mink farm operations (Shriner et al. 2021). Antibodies to SARS-CoV-2 were detected in white-tailed deer (*Odocoileus virginianus*) indicating exposure to the virus (Chandler et al. 2021), and infection was later confirmed in this species (Hale et al. 2022). Captive wild animals in zoos, particularly members of the Felidae family, were infected with the virus (McAloose et al. 2020), and several North American species have experimentally been shown susceptible to the virus, including deer mice (*Peromyscus maniculatus*), striped skunks (*Mephitis mephitis*), and bushy-tailed woodrats (*Neotoma cinerea*) (Bosco-Lauth et al. 2021).

Because evidence indicates that SARS-CoV-2 originated in Asian bats (Zhou et al. 2020; Zhou et al. 2021; Wacharapluesadee et al. 2021), concern that the virus could infect North American bat species has been raised, particularly for bat populations under severe threat from another pathogen, *Pseudogymnoascus destructans* (Cheng et al. 2021; Hoyt et al. 2021). Whether North American bats could provide a reservoir of the virus and additional routes of transmission to humans and other susceptible species, as well as any effects of SARS-CoV-2 on populations are important to determine, as are any management measures that could be used to help protect these populations.

Big brown bats (*Eptesicus fuscus*) were previously challenged with SARS-CoV-2 and demonstrated resistance to infection (Hall et al. 2021). This species often encounters humans, as they frequently reside in anthropogenic structures, including occupied homes and other buildings. Another common North American bat species, the Mexican free-tailed bat, (TABR: *Tadarida brasiliensis*), resides in very large colonies in the southern United States, often in urban areas. This species is migratory and if susceptible to SARS-CoV-2, could transport the virus to/from Central and South America on their migratory routes. In this study we challenged TABR with SARS-CoV-2 to determine their susceptibility to infection, reservoir potential, the adaptability of the virus to a new potential host, and potential effects of the virus on their populations.

## Materials and methods

### Virus acquisition and propagation

We obtained the SARS-CoV-2 virus (2019-nCoV/USA-WA1/2020) from BEI Resources, National Institute of Allergy and Infectious Diseases (NIAID), National Institutes of Health (NIH) (Manassas, Virginia)The virus was isolated from the first confirmed patient with coronavirus disease 2019 (COVID-19) in the United States (Harcourt et al. 2020). We propagated and quantified the virus in Vero E6 cell culture using standard techniques.

### Animal acquisition and husbandry

All husbandry and experimental protocols were approved by the U.S. Geological Survey (USGS) National Wildlife Health Center (NWHC) Institutional Animal Care and Use Committee. Wild TABR were captured in Williamson County, Texas in August 2021. A mixture of adult and juvenile male bats was collected and immediately placed into a temperature-controlled chamber maintained at approximately 20°C. This temperature induced the bats to enter torpor during transport to the NWHC, Madison, Wisconsin.

On arrival at the NWHC, the bats were given veterinary examinations and treated topically with selamectin for parasites (Zoetis, Florham Park, New Jersey). The bats were hand fed mealworms (*Tenebrio molito*) supplemented with a vitamin and mineral mixture, and water was provided *ad libitum*. Bats underwent a quarantine and acclimatization period of 30 days prior to commencement of this study during which time the bats learned to feed themselves.

### Pre-inoculation fecal sampling and coronavirus analysis

During the acclimatization period, we collected fecal samples from the individual bats to determine the presence of other coronaviruses in these subjects. Each fecal sample was suspended 10% (w/v) in viral transport medium (VTM; Hanks Balanced Salt Solution, 0.05% gelatin, 5% glycerin, 1500 units/ml penicillin, 1500 mg/ml streptomycin, 0.1 mg/ml gentamicin,1 mg/ml fungizone). Viral RNA was extracted using the MagMax Pathogen RNA/DNA kit (Applied Biosystems, Forest City, California) on a Kingfisher Flex magnetic particle processor according to the manufacturer’s instructions. The presence of coronaviruses was determined using methods previously described (Decaro and Larusso, 2020).

### Virus inoculation

Experimental inoculations were performed under Biosafety Level-3 conditions at the NWHC. We utilized 21 male Mexican free-tailed bats after the acclimatization period and pairs of bats were cohoused in mesh cages. One bat from each of nine bat pairs was inoculated with SARS-CoV-2, and its cagemate was left uninoculated to determine if the virus could be horizontally transmitted between bats. One bat was inoculated and housed individually. The SARS-CoV-2 inoculum dose was 10^5^ TCID_50_ /bat and was administered nasally (4 µl) and orally (6 µl) using a micropipette. One bat pair was sham inoculated with the same volume of VTM. This technique has been used to inoculate other species with SARS-CoV-2 (Munster et al. 2020; Schlottau et al. 2020; Shi et al. 2020; Hall et al. 2021). The inoculum titer was verified by quantitative reverse transcription-polymerase chain reaction (qRT-PCR) as described below and virus viability confirmed in cell culture using Vero E6 cells.

### Animal monitoring and sampling

Bats were observed at least twice daily to monitor health status and document development of clinical signs. Just prior to inoculation and every other day thereafter, each bat was weighed, and oropharyngeal and rectal swabs (Puritan Medical Products, Guilford, Maine) were collected and placed in 0.5 ml VTM. On day post-inoculation (DPI) 7 and on DPI 14, bats from one cage (one inoculated bat, one uninoculated) were euthanized, a postmortem examination was conducted, and tissues and blood collected. At the end of the study (DPI 20), all remaining bats were euthanized and postmortem examinations were completed for the control bats and an additional cage pair. Blood was collected for serological analyses from all euthanized bats.

### qRT-PCR analyses

RNA extractions of swab material were performed in 96-well plates using Mag Max-96 AI/ND Viral RNA Isolation Kit (Applied Biosystems, Foster City, California) following the manufacturer’s instructions. A positive control sample consisting of a 1:100 dilution of the SARS-CoV-2 inoculum used in the study was included with each extraction series to validate successful RNA extraction. qRT-PCR analyses were conducted utilizing the Centers for Disease Control 2019-nCoV N1 primers and probe (https://www.cdc.gov/coronavirus/2019-ncov/lab/rt-pcr-panel-primer-probes.html) and AgPath-ID One-Step RT-PCR reagents (Ambion/ThermoFisher, Waltham, Massachusetts). We included a standard curve of serial dilutions of RNA extracted from SARS-CoV-2 virus stock (10^7^ TCID50/ml) in each qRT-PCR assay to quantify viral amounts.

### Rabies Diagnostics

Brain tissue was assessed for rabies infection using the direct fluorescent antibody test (DFA). After brain impressions were fixed in acetone, slides were stained with a FITC-labelled monoclonal antibody (mAB) conjugate (Fujirebio U.S. Inc., Malvern, Pennsylvania, USA) and visualized under a fluorescent microscope (Dean et al.1996).

### Necropsy and histopathology

Two animals (inoculated and uninoculated cagemates) were euthanized at DPI 7 (bats 127, 128) and DPI 14 (bats 103, 104), and an additional 2 sets of cagemates at DPI 20 (bats 109, 110, 111, 117), using an overdose of isoflurane with subsequent decapitation. Two uninoculated control animals (102, 108) were also euthanized at DPI 20. These subjects were immediately necropsied after euthanasia and body condition and gross observations were recorded. Portions of the nares, caudal lung, cranial lung, heart, liver, spleen, kidney, small intestine, colon and brain were collected and saved frozen at -80°C for virological analyses. Additional tissue portions were fixed in 10% neutral buffered formalin for histological analysis. For histopathological examination, fixed tissues were processed routinely, sectioned at approximately 5 µm and stained with hematoxylin and eosin at the Wisconsin Veterinary Diagnostic Laboratory (Madison, Wisconsin). At DPI 20, all remaining bats were euthanized, serum was collected, and all bat carcasses were saved frozen. Three bats (118, 123, 124) that were saved frozen and later shown to be infected by swab analysis were subsequently necropsied and sampled.

### SARS-CoV-2-specific immunohistochemistry (IHC)

For IHC, 4 µm sections of formalin-fixed paraffin-embedded tissue were mounted on positively charged Superfrost® Plus slides and subjected to IHC using an anti-nucleocapsid rabbit monoclonal antibody (HL344, Cell Signaling Technology, Danvers, Massachusetts). IHC was performed using the automated BOND-RXm platform and the Polymer Refine Red Detection kit (Leica Biosystems, Wetzlar, Germany). Following automated deparaffinization, heat-induced epitope retrieval (HIER) was performed using a ready-to-use citrate-based solution (pH 6.0; Leica Biosystems) at 100°C for 20 min. Sections were then incubated with the primary antibody (diluted at 1:1,600 in primary antibody diluent [Leica Biosystems]) for 30 min at room temperature, followed by a polymer-labeled goat anti-rabbit IgG coupled with alkaline phosphatase (30 min). Fast Red was used as the chromogen (15 minutes), and counterstaining was performed with hematoxylin for 5 min. Slides were dried in a 60 °C oven for 30 min and mounted with a permanent mounting medium (Micromount®, Leica Biosystems). Lung sections from a SARS-CoV-2-infected hamster were used as positive assay controls.

### Virus RNA extraction and qRT-PCR from bat tissues

Approximately 10 mg of each tissue was macerated in extraction buffer and RNA extracted using the ZYMO Research Quick DNA/RNA Pathogen Miniprep kit (ZYMO Research, Irvine, California) according to the manufacturer’s directions. qRT-PCR analyses were performed as described above.

### Antibody detection

To detect neutralizing antibodies to SARS-CoV-2, bat sera were screened at a 1:10 dilution using a competitive enzyme linked immunosorbent assay (SARS-CoV-2 sVNT, GenScript, Piscataway, New Jersey) according to the manufacturer’s instructions. As directed, a reduction in optical density (OD) of ≥ 30% compared to the mean OD of the negative control was considered positive for the presence of neutralizing antibodies. In addition to the positive control provided in the kit, we used positive guinea pig serum obtained through BEI Resources, NIAID, NIH: Polyclonal Anti-SARS Coronavirus antiserum (Guinea Pig, NR-10361). To determine neutralizing antibody titers from positive sera, samples were two-fold serially diluted and titers recorded as the reciprocal of the end-point dilution where the serum was no longer considered positive.

### Virus recovery and whole genome sequencing

Oral swab VTM from Bat 118 DPI 8 was inoculated into Vero E6 cells and incubated at 37 °C, 5% CO2 for 7 days. The flasks were examined daily for cytopathic effects. Cell lysates collected after three cycles of freezing and thawing were used for RNA extraction and serial passage. Extracted RNA was converted to cDNA with SuperScript IV (ThermoFisher, Waltham, Massachusetts) or Maxima H minus (ThermoFisher, Waltham, Massachusetts) reverse transcriptase according to manufacturer’sinstructions. Tiled amplicon sequencing by the ARTIC method (Tyson et al. 2020) was performed using Oxford Nanopore Technology’s MinION running on a MK1C instrument. Bioinformatic analysis was performed using the CLC Genomics Workbench v22 (Qiagen, Redwood City, California) using a publicly available workflow (https://storage.googleapis.com/theiagen-resources/qiagen/SARS-CoV-2_Tutorial.zip).

## Results

### Presence of coronaviruses in Mexican free-tailed bats prior to inoculation

RT-PCR analyses of fecal material collected from the TABR prior to virus challenge revealed no evidence of alpha- or betacoronavirus infection, in any subject (Supplemental Table 1).

**Table 1.**
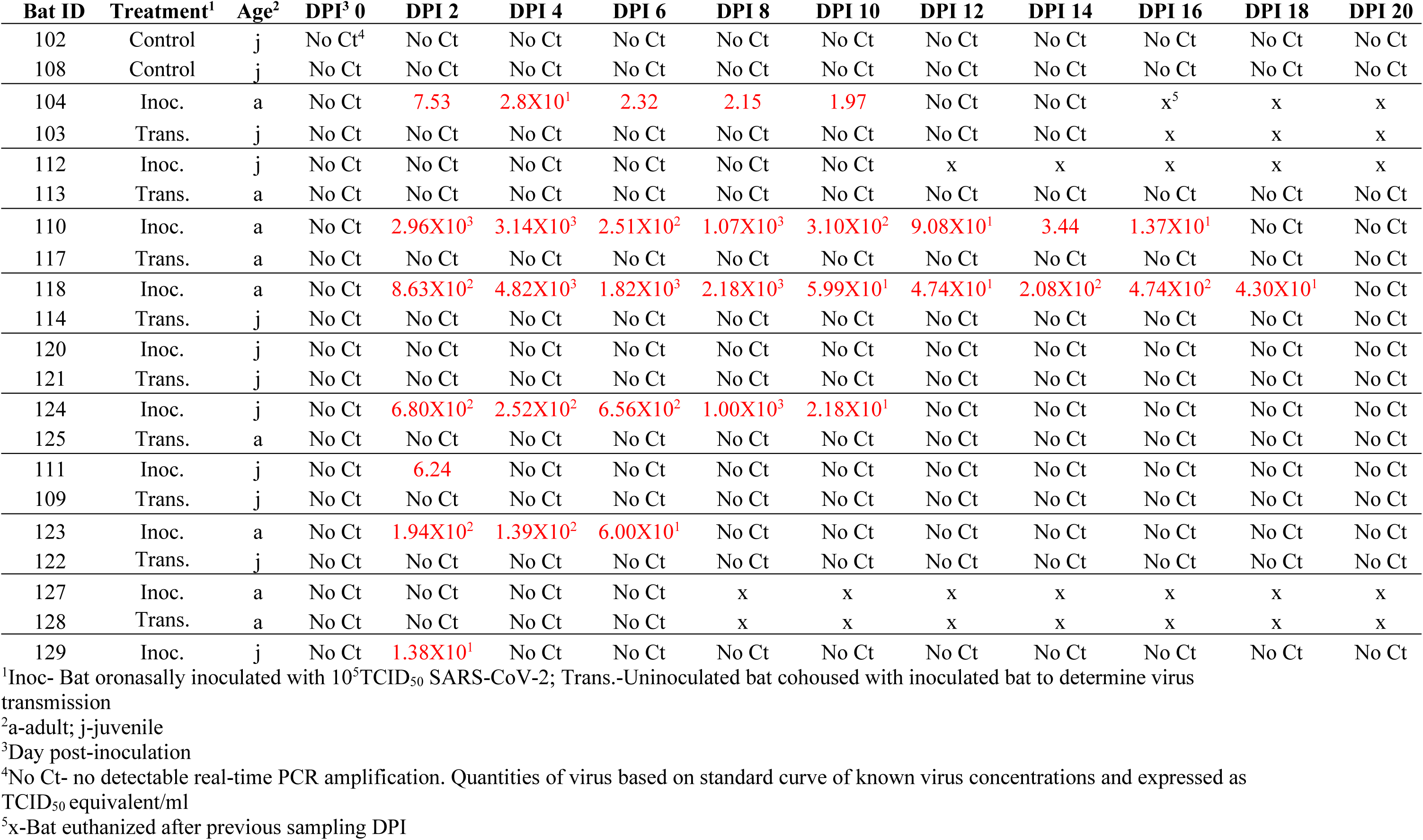
Quantitative RT-PCR analyses of oral swabs from Mexican free-tailed bats inoculated with SARS-CoV-2.

| Bat ID | Treatment<sup>1</sup> | Age<sup>2</sup> | DPI<sup>3</sup> 0 | DPI 2 | DPI 4 | DPI 6 | DPI 8 | DPI 10 | DPI 12 | DPI 14 | DPI 16 | DPI 18 | DPI 20 |
| --- | --- | --- | --- | --- | --- | --- | --- | --- | --- | --- | --- | --- | --- |
| 102 | Control | j | No Ct<sup>4</sup> | No Ct | No Ct | No Ct | No Ct | No Ct | No Ct | No Ct | No Ct | No Ct | No Ct |
| 108 | Control | j | No Ct | No Ct | No Ct | No Ct | No Ct | No Ct | No Ct | No Ct | No Ct | No Ct | No Ct |
| 104 | Inoc. | a | No Ct | 7.53 | 2.8X10<sup>1</sup> | 2.32 | 2.15 | 1.97 | No Ct | No Ct | x<sup>5</sup> | x | x |
| 103 | Trans. | j | No Ct | No Ct | No Ct | No Ct | No Ct | No Ct | No Ct | No Ct | x | x | x |
| 112 | Inoc. | j | No Ct | No Ct | No Ct | No Ct | No Ct | No Ct | x | x | x | x | x |
| 113 | Trans. | a | No Ct | No Ct | No Ct | No Ct | No Ct | No Ct | No Ct | No Ct | No Ct | No Ct | No Ct |
| 110 | Inoc. | a | No Ct | 2.96X10<sup>3</sup> | 3.14X10<sup>3</sup> | 2.51X10<sup>2</sup> | 1.07X10<sup>3</sup> | 3.10X10<sup>2</sup> | 9.08X10<sup>1</sup> | 3.44 | 1.37X10<sup>1</sup> | No Ct | No Ct |
| 117 | Trans. | a | No Ct | No Ct | No Ct | No Ct | No Ct | No Ct | No Ct | No Ct | No Ct | No Ct | No Ct |
| 118 | Inoc. | a | No Ct | 8.63X10<sup>2</sup> | 4.82X10<sup>3</sup> | 1.82X10<sup>3</sup> | 2.18X10<sup>3</sup> | 5.99X10<sup>1</sup> | 4.74X10<sup>1</sup> | 2.08X10<sup>2</sup> | 4.74X10<sup>2</sup> | 4.30X10<sup>1</sup> | No Ct |
| 114 | Trans. | j | No Ct | No Ct | No Ct | No Ct | No Ct | No Ct | No Ct | No Ct | No Ct | No Ct | No Ct |
| 120 | Inoc. | j | No Ct | No Ct | No Ct | No Ct | No Ct | No Ct | No Ct | No Ct | No Ct | No Ct | No Ct |
| 121 | Trans. | j | No Ct | No Ct | No Ct | No Ct | No Ct | No Ct | No Ct | No Ct | No Ct | No Ct | No Ct |
| 124 | Inoc. | j | No Ct | 6.80X10<sup>2</sup> | 2.52X10<sup>2</sup> | 6.56X10<sup>2</sup> | 1.00X10<sup>3</sup> | 2.18X10<sup>1</sup> | No Ct | No Ct | No Ct | No Ct | No Ct |
| 125 | Trans. | a | No Ct | No Ct | No Ct | No Ct | No Ct | No Ct | No Ct | No Ct | No Ct | No Ct | No Ct |
| 111 | Inoc. | j | No Ct | 6.24 | No Ct | No Ct | No Ct | No Ct | No Ct | No Ct | No Ct | No Ct | No Ct |
| 109 | Trans. | j | No Ct | No Ct | No Ct | No Ct | No Ct | No Ct | No Ct | No Ct | No Ct | No Ct | No Ct |
| 123 | Inoc. | a | No Ct | 1.94X10<sup>2</sup> | 1.39X10<sup>2</sup> | 6.00X10<sup>1</sup> | No Ct | No Ct | No Ct | No Ct | No Ct | No Ct | No Ct |
| 122 | Trans. | j | No Ct | No Ct | No Ct | No Ct | No Ct | No Ct | No Ct | No Ct | No Ct | No Ct | No Ct |
| 127 | Inoc. | a | No Ct | No Ct | No Ct | No Ct | x | x | x | x | x | x | x |
| 128 | Trans. | a | No Ct | No Ct | No Ct | No Ct | x | x | x | x | x | x | x |
| 129 | Inoc. | j | No Ct | 1.38X10<sup>1</sup> | No Ct | No Ct | No Ct | No Ct | No Ct | No Ct | No Ct | No Ct | No Ct |

<sup>1</sup>Inoc- Bat oronasally inoculated with 10<sup>5</sup>TCID<sub>50</sub> SARS-CoV-2; Trans.-Uninoculated bat cohoused with inoculated bat to determine virus transmission
<sup>2</sup>a-adult; j-juvenile
<sup>3</sup>Day post-inoculation
<sup>4</sup>No Ct- no detectable real-time PCR amplification. Quantities of virus based on standard curve of known virus concentrations and expressed as TCID<sub>50</sub> equivalent/ml
<sup>5</sup>x-Bat euthanized after previous sampling DPI

### Rabies virus infection

Prior to SARS-CoV-2 inoculation, one bat exhibited loss of appetite, aggressive behavior towards its cagemate and weight loss over 3-4 days. We euthanized this bat and tested for rabies. It was positive for the presence of rabies virus by DFA, therefore we tested all the remaining TABR after euthanasia and all were negative for the presence of rabies virus (Supplemental Table 1).

### SARS-CoV-2 excretion after experimental inoculation

qRT-PCR analysis of oral swabs is shown in Table 1. Of the 10 SARS-CoV-2 inoculated TABR, five (Bats 104, 110, 118, 124, 123) excreted detectable viral RNA. Two additional bats (111, 129) had high cycle threshold (Ct) readings only on the first sampling after inoculation (DPI 2) and likely was detection of residual inoculum. The duration of oral excretion ranged from DPI 6 (Bat 123), up to 18 DPI (Bat 118). The maximum amount of virus detected on an oral swab was 4.82×10^3^ TCID50 equivalent/ml (bat 118, DPI 4). In contrast to oral excretion, no virus was detected in rectal swabs from any TABR (Table 2). The effect of the bat’s age on susceptibility was inconclusive as four of the infected bats were adults and one was a juvenile, but the numbers were too small to be reliable.

**Table 2.** Quantitative RT-PCR analyses of rectal swabs from Mexican free-tailed bats inoculated with SARS-CoV-2.

| Bat ID | Treatment<sup>1</sup> | DPI<sup>2</sup> 0 | DPI 2 | DPI 4 | DPI 6 | DPI 8 | DPI 10 | DPI 12 | DPI 14 | DPI 16 | DPI 18 | DPI 20 |
| --- | --- | --- | --- | --- | --- | --- | --- | --- | --- | --- | --- | --- |
| 102 | Control | No Ct<sup>3</sup> | No Ct | No Ct | No Ct | No Ct | No Ct | No Ct | No Ct | No Ct | No Ct | No Ct |
| 108 | Control | No Ct | No Ct | No Ct | No Ct | No Ct | No Ct | No Ct | No Ct | No Ct | No Ct | No Ct |
| 104 | Inoc. | No Ct | No Ct | No Ct | No Ct | No Ct | No Ct | No Ct | No Ct | x<sup>4</sup> | x | x |
| 103 | Trans. | No Ct | No Ct | No Ct | No Ct | No Ct | No Ct | No Ct | No Ct | x | x | x |
| 112 | Inoc. | No Ct | No Ct | No Ct | No Ct | No Ct | No Ct | x | x | x | x | x |
| 113 | Trans. | No Ct | No Ct | No Ct | No Ct | No Ct | No Ct | No Ct | No Ct | No Ct | No Ct | No Ct |
| 110 | Inoc. | No Ct | No Ct | No Ct | No Ct | No Ct | No Ct | No Ct | No Ct | No Ct | No Ct | No Ct |
| 117 | Trans. | No Ct | No Ct | No Ct | No Ct | No Ct | No Ct | No Ct | No Ct | No Ct | No Ct | No Ct |
| 118 | Inoc. | No Ct | No Ct | No Ct | No Ct | No Ct | No Ct | No Ct | No Ct | No Ct | No Ct | No Ct |
| 114 | Trans. | No Ct | No Ct | No Ct | No Ct | No Ct | No Ct | No Ct | No Ct | No Ct | No Ct | No Ct |
| 120 | Inoc. | No Ct | No Ct | No Ct | No Ct | No Ct | No Ct | No Ct | No Ct | No Ct | No Ct | No Ct |
| 121 | Trans. | No Ct | No Ct | No Ct | No Ct | No Ct | No Ct | No Ct | No Ct | No Ct | No Ct | No Ct |
| 124 | Inoc. | No Ct | No Ct | No Ct | No Ct | No Ct | No Ct | No Ct | No Ct | No Ct | No Ct | No Ct |
| 125 | Trans. | No Ct | No Ct | No Ct | No Ct | No Ct | No Ct | No Ct | No Ct | No Ct | No Ct | No Ct |
| 111 | Inoc. | No Ct | No Ct | No Ct | No Ct | No Ct | No Ct | No Ct | No Ct | No Ct | No Ct | No Ct |
| 109 | Trans. | No Ct | No Ct | No Ct | No Ct | No Ct | No Ct | No Ct | No Ct | No Ct | No Ct | No Ct |
| 123 | Inoc. | No Ct | No Ct | No Ct | No Ct | No Ct | No Ct | No Ct | No Ct | No Ct | No Ct | No Ct |
| 122 | Trans. | No Ct | No Ct | No Ct | No Ct | No Ct | No Ct | No Ct | No Ct | No Ct | No Ct | No Ct |
| 127 | Inoc. | No Ct | No Ct | No Ct | No Ct | x | x | x | x | x | x | x |
| 128 | Trans. | No Ct | No Ct | No Ct | No Ct | x | x | x | x | x | x | x |
| 129 | Inoc. | No Ct | No Ct | No Ct | No Ct | No Ct | No Ct | No Ct | No Ct | No Ct | No Ct | No Ct |

<sup>1</sup>Inoc- Bat oronasally inoculated with 10<sup>5</sup>TCID<sub>50</sub> SARS-CoV-2; Trans.-Uninoculated bat cohoused with inoculated bat to determine virus transmission
<sup>2</sup>Day post-inoculation
<sup>3</sup>No Ct- no detectable real-time PCR amplification. Quantities of virus based on standard curve of known virus concentrations and expressed as TCID<sub>50</sub> equivalents/ml.
<sup>4</sup>x-Bat euthanized after previous sampling DPI

An oral swab from a positive bat (Bat 118 DPI 8) was inoculated into Vero E6 tissue culture to verify viral viability. This resulted in positive virus isolates from the first and third passage that were confirmed to be SARS-CoV-2 by qRT-PCR analyses as described above. Whole genome sequencing of these isolates revealed that they shared the same genetic changes as the WA-1 isolate used as inoculum, when compared to the original Wuhan SARS-CoV-2 isolate. Two additional, consistent changes in both passages of the Bat 118 DPI 8 isolate were found in noncoding regions that were not in the WA-1 inoculum isolate. These are shown in Supplemental Table 2.

### Serological analyses

SARS-CoV-2 antibodies were detected by competitive Enzyme Linked Immunosorbent Assay (cELISA) in five TABR with percent inhibitions ranging from about 34 – 79% (Table 3). These five bats were the same five that were orally excreting virus detected by qRT-PCR (Table 1). These seropositive bat sera were subsequently tested at dilutions from 1:20 – 1:80. One serum (110) was positive only at the original 1:10 dilution. Two sera (123, 124) were positive at 1:20 whereas two sera (104, 118) remained positive at 1:40 dilutions.

**Table 3.** Competitive ELISA^1^ results of Mexican free-tailed bat sera tested after experimental challenge with SARS-CoV-2.

| Bat ID | % Inhibition<sup>2</sup> | Results<sup>3</sup> |  |  |  |
| --- | --- | --- | --- | --- | --- |
|  |  | 1:10 | 1:20 | 1:40 | 1:80 |
| 102* | -2.656 | Neg |  |  |  |
| 103 | 14.73 | Neg |  |  |  |
| 104 | 62.75 | Pos | Pos | Pos | Neg |
| 105<sup>4</sup> | -157.1 | Neg |  |  |  |
| 108* | 12.66 | Neg |  |  |  |
| 109 | 13.49 | Neg |  |  |  |
| 110 | 34.32 | Pos | Neg | Neg | Neg |
| 111 | 1.690 | Neg |  |  |  |
| 112 | 25.28 | Neg |  |  |  |
| 113 | 15.21 | Neg |  |  |  |
| 114 | 34.25 | Neg |  |  |  |
| 117 | 17.42 | Neg |  |  |  |
| 118 | 79.30 | Pos | Pos | Pos | Neg |
| 119<sup>4</sup> | 14.45 | Neg |  |  |  |
| 120 | 0.862 | Neg |  |  |  |
| 121 | 14.25 | Neg |  |  |  |
| 122 | -24.39 | Neg |  |  |  |
| 123 | 65.44 | Pos | Pos | Neg | Neg |
| 124 | 57.30 | Pos | Pos | Neg | Neg |
| 125 | -31.63 | Neg |  |  |  |
| 127 | -15.07 | Neg |  |  |  |
| 128 | 28.04 | Neg |  |  |  |
| 129 | 7.899 | Neg |  |  |  |

<sup>1</sup>SARS-CoV-2 sVNT, GenScript, Piscataway, New Jersey
<sup>2</sup>Inhibition of >30% is positive.
<sup>3</sup>Two-fold serial dilutions of positive sera
<sup>4</sup>Bats euthanized prior to the study
\*Uninoculated controls

### Clinical signs of infection, postmortem examination and histopathology

Over the three-week course of this study, no overt clinical signs of SARS-CoV-2 disease were observed in 18/21 bats, including the five bats that became infected and were orally excreting virus. These bats maintained or gained weight during the study (Table 4) and appeared healthy. One inoculated bat (127) presented with lethargy, decreased respiratory rate, and increased respiratory effort; and was euthanized on DPI 6, along with its cagemate (128). An additional bat (112) presented with lethargy, obtundation, weight loss, hypersalivation, inability to swallow, and respiratory distress; and was euthanized at DPI 10.

**Table 4.** Body weights (g) of SARS-CoV-2 inoculated Mexican free-tailed bats

| **<u>Bat ID</u>** | **<u>Treatment</u>** | **<u>DPI<sup>1</sup> 0</u>** | **<u>DPI 2</u>** | **<u>DPI 4</u>** | **<u>DPI 6</u>** | **<u>DPI 8</u>** | **<u>DPI 10</u>** | **<u>DPI 12</u>** | **<u>DPI 14</u>** | **<u>DPI 16</u>** | **<u>DPI 18</u>** | **<u>DPI 20</u>** |
| --- | --- | --- | --- | --- | --- | --- | --- | --- | --- | --- | --- | --- |
| 102 | Control | 10.19 | 10.11 | 11.23 | 11.27 | 11.8 | 12.13 | 11.78 | 11.7 | 13.45 | 13.13 | 13.24 |
| 108 | Control | 11.97 | 11.42 | 12.1 | 11.86 | 12.34 | 12.48 | 12.3 | 12.93 | 12.31 | 14.38 | 14.46 |
| 104 | Inoculated | 11.99 | 10.56 | 11.85 | 12.13 | 12.9 | 12.76 | 12.81 | 13.11 | x<sup>2</sup> | x | x |
| 103 | Transmission | 11.81 | 10.31 | 11.3 | 11.28 | 11.56 | 12.12 | 11.88 | 12.81 | x | x | x |
| 112 | Inoculated | 11.75 | 11.97 | 11.31 | 9.72 | 9.26 | 8.4 | x | x | x | x | x |
| 113 | Transmission | 14.37 | 13.25 | 12.84 | 13.81 | 14.13 | 15.08 | 14.83 | 15.01 | 14.67 | 14.43 | 14.43 |
| 110 | Inoculated | 16.19 | 14.34 | 15.39 | 15.14 | 15.36 | 15.6 | 16.03 | 15.09 | 16.04 | 16.46 | 16.66 |
| 117 | Transmission | 12.45 | 11.26 | 11.84 | 11.53 | 11.83 | 12.87 | 12.86 | 12.18 | 12.66 | 13.09 | 13.13 |
| 118 | Inoculated | 13.45 | 12.34 | 12.87 | 12.56 | 12.9 | 13.57 | 13.7 | 13.72 | 13.84 | 13.91 | 13.75 |
| 114 | Transmission | 10.85 | 10.2 | 10.92 | 10.66 | 11.31 | 11.61 | 11.7 | 11.39 | 11.29 | 11.27 | 13.68 |
| 120 | Inoculated | 12.1 | 10.73 | 11.71 | 11.5 | 11.63 | 12.14 | 12.27 | 12.48 | 12.9 | 12.47 | 12.37 |
| 121 | Transmission | 10.33 | 11.1 | 12 | 11.53 | 11.55 | 12.05 | 11.97 | 12.04 | 12.54 | 12.49 | 13.11 |
| 124 | Inoculated | 13.25 | 11.95 | 12.87 | 12.52 | 13.33 | 13.73 | 14.05 | 14.49 | 14.44 | 14.09 | 14.9 |
| 125 | Transmission | 13.07 | 12.2 | 12.45 | 12.16 | 11.57 | 12.05 | 12.19 | 12.41 | 12.36 | 12.56 | 12.84 |
| 111 | Inoculated | 14.02 | 13.11 | 13.15 | 13.73 | 13.62 | 13.92 | 14.25 | 13.95 | 13.98 | 13.64 | 13.93 |
| 109 | Transmission | 10.86 | 11.29 | 11.14 | 10.56 | 10.64 | 10.99 | 11.39 | 12.05 | 12.14 | 11.97 | 12.23 |
| 123 | Inoculated | 11.09 | 10.45 | 10.83 | 10.31 | 10.62 | 11.19 | 11.5 | 11.78 | 11.48 | 11.84 | 12.12 |
| 122 | Transmission | 9.23 | 9.45 | 9.78 | 10.33 | 10.6 | 11.37 | 11.72 | 11.57 | 11.65 | 11.53 | 11.71 |
| 127 | Inoculated | 9.13 | 8.86 | 9.37 | 8.69 | x | x | x | x | x | x | x |
| 128 | Transmission | 8.39 | 8.12 | 8.07 | 7.95 | x | x | x | x | x | x | x |
| 129 | Inoculated | 10.4 | 10.96 | 10 | 11.51 | 10.92 | 10.7 | 11.48 | 12.02 | 10.74 | 10.76 | 11.39 |

<sup>1</sup>Day post-inoculation493 <sup>2</sup>x-Bat euthanized after previous sampling DPI

Upon further examination, the bat that presented with respiratory difficulty (Bat 127) and its cagemate (128) were in poor or emaciated body condition, respectively, with minimal or no fat stores. Bat 112, euthanized due to respiratory distress and hypersalivation, was in fair body condition. All other bats were in good body condition evidenced by moderate to abundant fat stores. Gross findings included clear nasal discharge (112, 127), red foci or mottling in the lungs (all cases except 127), pulmonary congestion (102-104, 008-011, 117, 118, 128), intestinal pallor (102, 108, 109, 111, 117, 123, 124, 128), small foci of depigmentation of the patagia (103, 108, 111, 118, 123, 124, 127) and patagial tears (112, 128).

Histopathologic findings included pulmonary congestion, hemorrhage and alveolar collapse (all non-frozen cases), meningeal hemorrhage (103, 104, 111, 112, 127), pulmonary vascular thrombosis (112), large numbers of bacterial rods in bronchial epithelium and cardiac valve and parenchyma (127), pale cardiomyofibers (110, 112, 127), nasal cavity hemorrhage (102, 104, 109, 110, 111, 117, 128), necroulcerative and suppurative pharyngitis with intralesional bacteria (112), exocytosis of neutrophils in nasal turbinate epithelium (104), non-suppurative dermatitis (102, 108, 109, 110, 117), non-suppurative dermal myositis (102), mononuclear cells in hepatic sinusoids (108, 110, 111, 128), trematodes in the lumen of the bile ducts (110, 118) or gall bladder (118, 127), bile duct hyperplasia (110, 118), intestinal nematodiasis (104, 118, 127), intestinal cestodiasis (117, 127), intestinal trematodiasis (118), and intestinal luminal hemorrhage (127). The three additional bats (118, 123, 124) that excreted virus were thawed, necropsies were performed, and tissues were collected. The histologic evaluation of the lungs in these cases was severely hindered by freeze artifact.

These histopathologic findings observed for all examined bats were consistent with the euthanasia procedures utilized, parasitism, or bacterial infections, and were not consistent with findings observed in other animals experimentally infected with SARS-CoV-2 (Munster et al. 2020; Schlottau et al. 2020; Shi et al. 2020).

### Immunohistochemistry for detection of SARS-CoV-2 antigen

Sections from the rostral nasal cavity, lung, heart, spleen, liver, pancreas, stomach, small and large intestine, and brain from a total of 14 experimentally and mock inoculated bats were subjected to immunohistochemistry for the detection of SARS-CoV-2 antigen. No viral antigen was detected in any of the tissues examined from these bats (Supplemental Table 3).

### SARS-CoV-2 in bat tissues

qRT-PCR analyses of tissues collected from the infected and control bats did not detect viral RNA in any tissue from any of the bats examined (Table 5).

**Table 5.** qRT-PCR analyses of selected tissues from SARS-CoV-2 infected and uninoculated control Mexican free-tailed bats

| <u>Tissue</u> | <u>Bat ID</u> |  |  |  |  |  |  |
| --- | --- | --- | --- | --- | --- | --- | --- |
|  | <u>102<sup>1</sup></u> | <u>108<sup>1</sup></u> | <u>104</u> | <u>110</u> | <u>118</u> | <u>123</u> | <u>124</u> |
| Brain | NoCt<sup>2</sup> | NoCt | NoCt | NoCt | NoCt | NoCt | NoCt |
| Nares | NoCt | NoCt | NoCt | NoCt | NoCt | NoCt | NoCt |
| Lung cranial | NoCt | NoCt | NoCt | NoCt | NoCt | NoCt | NoCt |
| Lung, caudal | NoCt | NoCt | NoCt | NoCt | NoCt | NoCt | NoCt |
| Heart | NoCt | NoCt | NoCt | NoCt | NoCt | NoCt | NoCt |
| Liver | NoCt | NoCt | NoCt | NoCt | NoCt | NoCt | NoCt |
| Kidney | NoCt | NoCt | NoCt | NoCt | NoCt | NoCt | NoCt |
| Spleen | NoCt | NoCt | NoCt | NoCt | NoCt | NoCt | NoCt |
| Small intestine | NoCt | NoCt | NoCt | NoCt | NoCt | NoCt | NoCt |
| Colon | NoCt | NoCt | NoCt | NoCt | NoCt | NoCt | NoCt |

<sup>1</sup>Uninoculated control
<sup>2</sup>NoCt- no detectable RT-PCR reaction
Tissues collected from bats euthanized and necropsied on DPI 20

## Discussion

We experimentally challenged Mexican free-tailed bats with SARS-CoV-2. Of the ten inoculated bats, five orally excreted virus for 6-18 days post challenge. These five also mounted an immune response and cleared the virus before the end of the study. Based on these findings we concluded that Mexican free-tailed bats are susceptible to infection by SARS-CoV-2.

We found no evidence of transmission between cohoused TABR. Five of the ten (50%) inoculated bats became infected indicating that our inoculum titer (10^5^ TCID_50_/ dose) was apparently near the 50% infectious dose for this species. However, the largest amount of virus excreted by infected bats was between 10^3^ and 10^4^ TCID50 equivalents/ml. Thus, based on these data, contact transmission between TABR would be unlikely.

Another possible reason for no transmission between TABR was the way we housed the bats during the challenge study. In the wild, TABR roost in very dense colonies that aggregate in large numbers in natural and anthropogenic structures such as bridges, caves, and culverts. In our study we cohoused two bats, one inoculated and one uninoculated, in cages of approximately 1 meter^3^. This relatively large space did not force the bats to congregate as closely as occurs in natural roosts and thus “social distancing” may have impeded viral transmission. In future challenge studies we plan to adjust the bat housing to help take this factor into account.

In addition to the apparent lack of transmission, another observation from this study is that the virus was not excreted via the digestive system. Given the sizes of TABR colonies, large amounts of fecal material accumulate that could potentially become a source of infectious virus. We found no evidence of virus in the digestive tracts or in rectal swabs of any bat, including the infected bats.

Another finding was that TABR showed no obvious adverse health effects from SARS-CoV-2 infection. While it is unknown if infected wild bats would have diminished capacity to forage for food, perform maternal care, or other life functions, our findings indicate that TABR populations are likely not at risk from the pandemic.

Regardless, an accurate determination of the infectious dose of SARS-CoV-2 in TABR would be an important next study. We do not know if the amounts of virus excreted are enough to infect other mammalian species, including humans or if TABR can be infected with SARS-CoV-2 by exposure to sick humans. Because this species often resides in urban settings, these are important public health and pandemic ecological concerns.

We were unable to conduct necropsies and collect tissues from actively infected animals. All bats euthanized for these purposes during the study were not among the infected cohort, and by the end of the study, all infected bats had cleared the virus. Therefore, the qRT-PCR, pathology and histochemistry results from bat tissues were inconclusive.

It is generally accepted that SARS-CoV-2 originated in wild horseshoe bats (*Rhinolophus sp*.) from China, subsequently transmitted to other host species, and ultimately infected humans, leading to a pandemic (Zhou et al. 2020). The virus also infects at least one other bat species, Egyptian fruit bats, *Rousettus aegyptiacus* (Schlottau et al. 2020). This has raised concerns that the virus could infect North American bats, some of whose populations are under severe pressure from other diseases and from habitat degradation. Hall et al. 2021 previously showed that big brown bats, a species commonly encountered by humans, is resistant to infection by SARS-CoV-2. In this study, however, we demonstrated that TABR, a migratory bat that resides in large colonies, often in urban areas, is susceptible to infection by this virus. Based on the comparative structure of the SARS-CoV-2 cellular receptor, Damas et al. (2020) predicted that TABR were unlikely to become infected with SARS-CoV-2. Thus, it is likely that other factors are involved in mediating susceptibility to this virus. The genetic changes we detected in viral isolates from the swab of one infected bat indicates that mutations can occur rapidly in a new host and that genetic analyses of recovered viral isolates may further inform our understanding of the emergence of novel viral variants. Our findings also indicate that susceptibility of each species to SARS-CoV-2 is independent, and each species would benefit from being examined individually. These results have implications for bat rehabilitators, wildlife biologists, cave recreationists, and the public at large, if they contact Mexican free-tailed bats or enter caves or other environments where bats are roosting.

Supporting data are available in Hall et al. (2022) https://doi.org/10.5066/P9RDA1H6.

## Acknowledgements

This work was funded by the U.S. Geological Survey’s Ecosystems Management Area.

M. Carossino received support from the Center for Lung Biology and Disease (CLBD), Center of Biomedical Research Excellence, National Institute of General Medical Sciences of the National Institutes of Health under P20GM130555. We thank Katy Griffin, Jeffrey Messer, Harrison Lamb, Lauren Dycee-Holtz, Rachel Lambert, Carrie Allison Smith, and Casey Hall, for technical contributions. We also thank members of the histopathology and immunohistochemistry section at the Louisiana Animal Disease Diagnostic Laboratory for their assistance. We are particularly indebted to the NWHC Animal Services staff and volunteers, without whose assistance, diligence and hard work this study could not have been accomplished. We are also indebted to the Texas Parks and Wildlife Department for coordinating live bat acquisition. The use of trade, firm, or product names is for descriptive purposes only and does not imply endorsement by the U.S. Government. The authors acknowledge they have no personal financial interests or conflicts of interest with this research article.

---

**Supplemental Table 1.**
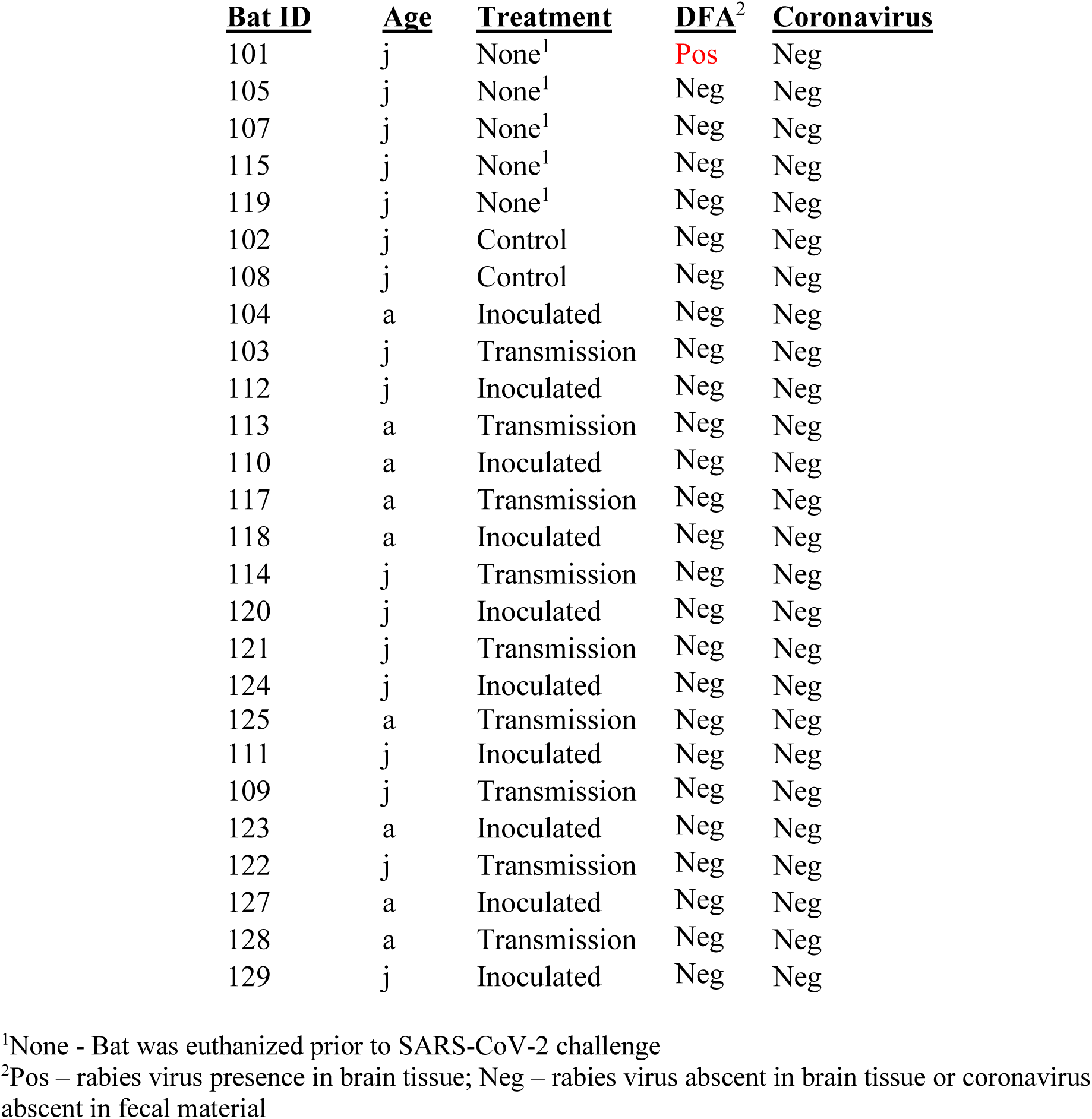
Rabies diagnostic results from Mexican free-tailed bats by direct fluorescent antibody (DFA) and the presence of alpha-or betacoronaviruses prior to initiation of the SARS-CoV-2 challenge study.

**Supplemental Table 2.**
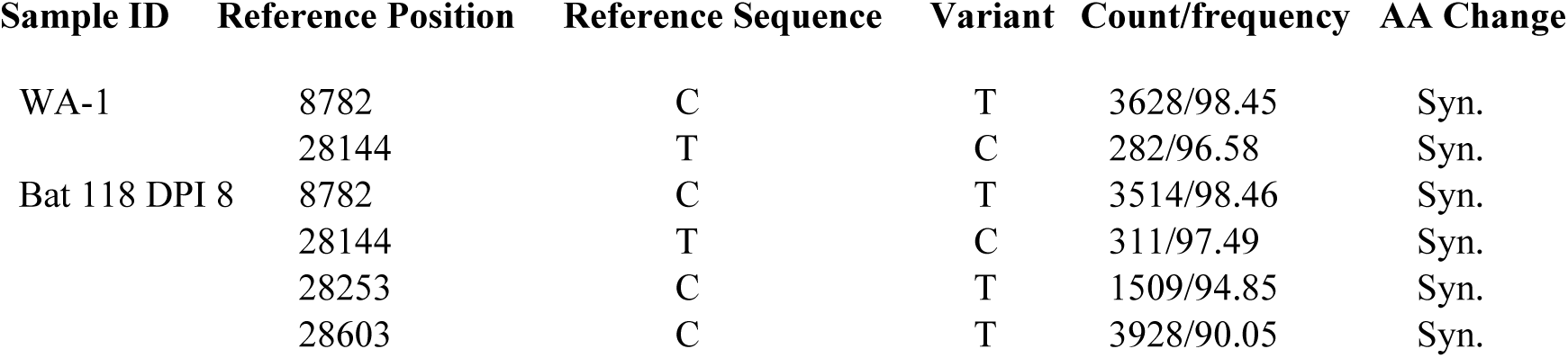
Variant detection in SARS-CoV-2 isolated from Bat 118, DPI 8. Genetic changes in the SARS-CoV-2 inoculum strain (USA-WA1/2020; Genbank MN985325) and Bat 118 DPI 8 (Genbank OM995890) after one passage in Vero cells compared with the reference strain (Wuhan HU-1; Genbank MN908947). Sample ID-sample name. Reference position-location of genetic variant on Wuhan HU-1 genome. Reference sequence-nucleotide sequence in HU-1. Variant-nucleotide in WA-1 and bat isolate. Count-number of reads covering the variant. Frequency-proportions of reads with variant. AA change-change in amino acid coding affected by variant. Syn.- synonymous

**Supplemental Table 3.**
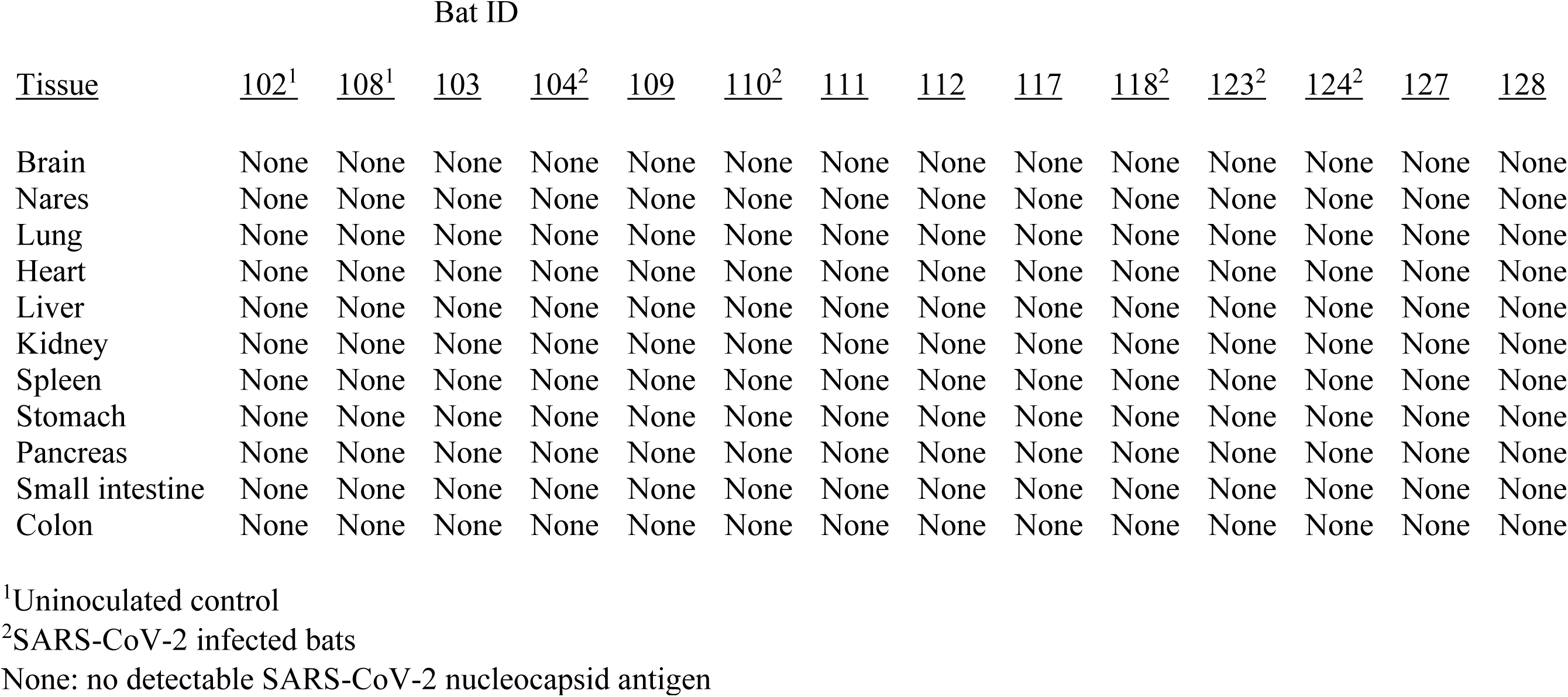
SARS-CoV-2 antigen distribution in selected tissues from SARS-CoV-2 inoculated and uninoculated control Mexican free-tailed bats. Bats euthanized, necropsied and tissue samples collected on day post-inoculation (DPI) 7 (bats 127, 128)); DPI 10 (bat 112); DPI 14 (bats 103, 104); and DPI 20 for the remaining bats.

